# A novel workflow integrating whole-body PET microdosing data and therapeutic-dose pharmacokinetics across species to inform first-in-human dose selection

**DOI:** 10.64898/2026.09.03.748870

**Authors:** Bach Tran Nguyen, Aurélie Barrail-Tran, Moreno Ursino, Sébastien Goutal, Fabien Caillé, Thibaut Naninck, Roger Le Grand, Olivier Lambotte, Nicolas Tournier, Emmanuelle Comets

## Abstract

**Introduction:** First-in-human studies require dose extrapolation from pharmacokinetic animal studies combined with safety assessments. Subtherapeutic doses of radiolabelled drugs can be administered in preclinical and early clinical development to gain a dynamic pharmacokinetic understanding, potentially informing pharmacologically-based pharmacokinetics (PBPK) models through whole-body PET data.

**Aims:** to develop a PET-informed modelling framework for preclinical-to-human extrapolation, with dolutegravir as a case-study.

**Methods:** We developed a structured workflow integrating micro- and conventional-dose data for interspecies and dose extrapolation. We built a whole-body PBPK model in PK-Sim/MoBi (v12.1) using non-human primate (NHP) data in 5 key organs incorporating dolutegravir physico-chemical properties, protein binding, metabolism and efflux. Sensitivity analyses and parameter estimation were performed sequentially first with PET microdosing organ data over 3h, then with fluid and tissue concentrations following a 2.5 mg/kg IV injection. Finally, 100 Caucasian healthy adults (50% male, 20–80 years) receiving 50 mg qd po after high-fat meals were simulated using the two sets of estimated parameters and physiology-related parameter distributions provided by PK-Sim.

**Results:** Dolutegravir blood data were well described in NHPs over 3 hours, with parameters adjusted to handle the macrodose-related changes. While microdose-based parameter estimates systematically underpredicted exposure, combining NHP micro- and conventional dose data predicted steady-state geometric mean AUC_0-24_ and C_max_ closely matching human reported profiles, although slightly underpredicting C_trough_.

**Conclusions:** PET-PBPK modelling combining micro- and conventional doses in NHPs successfully predicted dolutegravir concentrations in healthy volunteers, additionally informing tissue distribution. This proof-of-concept study supports early PET data acquisition to build robust priors for first-in-human studies.

**Key Points:**

- This study presents the first integration of PET microdosing data from nonhuman primates into a whole-body physiologically based pharmacokinetic model, enabling quantitative extrapolation of dolutegravir pharmacokinetics to humans.
- The developed model using PET data and further calibrated using traditional plasma concentration profiles in nonhuman primates was able to predict human steady-state plasma exposure metrics (C_max_, AUC_0-24h_, half-life) in human.
- PET-informed modelling provides dynamic organ-specific tissue distribution, offering novel insights into drug disposition not feasible with conventional preclinical methods.
- This workflow holds clinical relevance by complementing phase I studies for refined first-in-human dose selection and early tissue exposure assessment, potentially accelerating safer drug development and limiting animal experiments.

## 1. INTRODUCTION

Drug development is often associated with high failure rates, especially at early proof-of-concept stages, with a key contributor being the limited predictive value of preclinical in vivo and in vitro models for extrapolating human pharmacokinetics (PK) and pharmacodynamics [1,2]. Additionally, standard phase I trials are long, costly and provide only restricted insight into tissue distribution and mechanistic biomarkers in the target population [2,3].

To address these translational gaps, regulatory frameworks from the FDA and ICH have formalised exploratory “phase 0” or microdosing studies, in which subtherapeutic doses are administered with reduced non-clinical requirements and no therapeutic intent, primarily to generate early data on PK, target engagement and mechanism of action. One of the prototypical technical implementations of a phase 0 (exploratory) clinical trial, PET microdosing approaches offers non-invasive, whole-body, high temporal-resolution characterisation of drug distribution, target engagement and early pharmacodynamic biomarkers at subtherapeutic exposures, thereby enabling first-in-human investigations with a substantial safety margin [2,4–7].

Physiologically based pharmacokinetic (PBPK) modelling offers a mechanistic means to integrate physicochemical, in vitro and preclinical in vivo data, most commonly plasma concentrations and less frequently PET imaging data. In this work, we propose a framework based on PBPK modelling to extrapolate from microdoses to therapeutic doses and from animals to human. Through this framework, we explore nonlinearities such as saturable ADME, and simulate human concentration–time profiles to inform dose selection in first-in-human studies. We showcase this approach using animal data from dolutegravir, a widely prescribed antiretroviral drug, which was given to NHPs either as a radiolabelled microdose followed by PET acquisition or as therapeutic doses followed by plasma and tissue concentration samples [8–10].

In the following, we first conceptualise the framework, then show the different steps of its application to dolutegravir: (i) develop a PBPK model of dolutegravir in non-human primates (NHPs) using sequentially PET microdosing and conventional PK data, (ii) perform simulation to extrapolate dolutegravir PK at clinically relevant doses in humans, (iii) compare the simulated concentration–time profiles to kinetics in healthy volunteers using literature data.

## 2. METHODS

### 2.1. PET-Informed Modelling Framework

In this study, we propose a novel PET-informed modelling framework (**Figure 1**) integrating dynamic microdosing PET and conventional PK data to predict therapeutic-dose profiles in humans. The framework is structured around different input datasets, with the objective of seamlessly integrating all available information for first-in-human predictions. We first present the generic framework, defining data, challenges and methodological approaches for the analysis and extrapolation, and in section 2.2, we showcase an application of the framework using available data for the dolutegravir case study including only animal data, and evaluate the framework by comparing the prediction of human PK to literature data.

**Figure 1.**
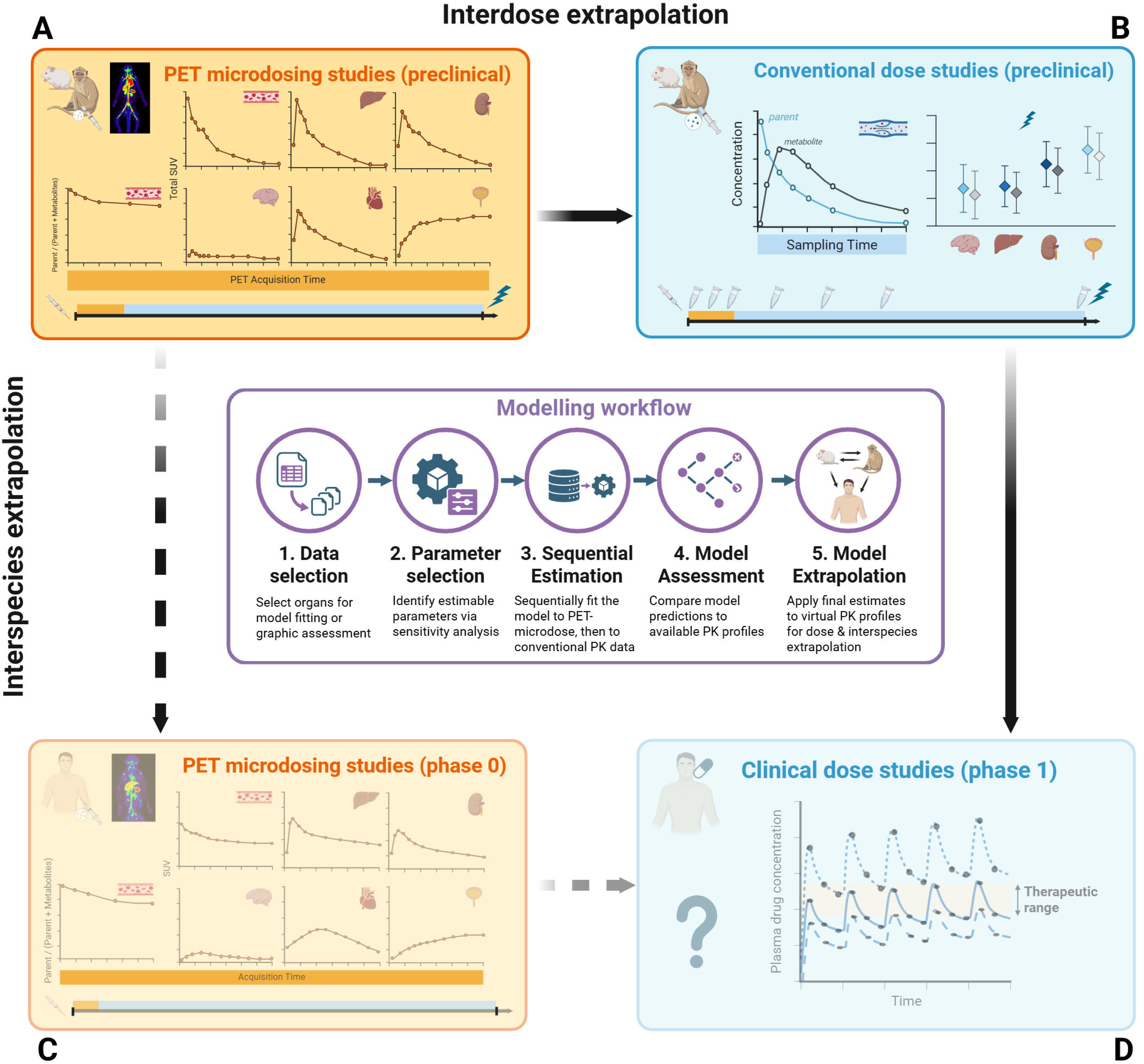
PET-informed modelling framework for human dose extrapolation.

#### 2.1.1. Modelling preclinical microdosing PET data (Step A)

PET imaging measures total radioactivity concentrations (parent drug and metabolites) over time in organs after the microdose injection of radio-labelled drugs. These studies therefore provide whole-body, dynamic data on drug distribution at sub-therapeutic doses (<1/100th of No-Observed Adverse Effect Level - NOAEL) [2,11]. These are constrained by the ability to radiolabel the drug of interest, the need for specialised radiochemistry and imaging infrastructure, due to onsite radiosynthesis of the radiolabelled compound prior to administration. The short acquisition windows (typically 1–3 h), dictated by radionuclide half-lives, limits the observation of slower elimination pathways. Moreover, PET data should be complemented with metabolite correction, e.g., via arterial plasma sampling and high-performance liquid chromatography (HPLC) quantification, to derive parent-specific plasma kinetics for comparison with image-derived time-activity curves (TACs) observed in organs.

These TACs can be used directly in PBPK models (as fraction of injected dose per organ volume) or converted to concentrations (using delineated organ volumes and injected mass). PBPK modelling is used at the core of our framework to leverage mechanistic models with predefined structure and parameter values obtained from in vitro assays, physicochemical properties, and species libraries. Because these models involve hundreds of parameters, a first step is to choose which parameters to estimate based on drug properties (lipophilicity, plasma protein binding, etc.) and physiology (known biological variability in renal/biliary clearance, enzyme/transporter V_max_, organ-dependant interstitial-plasma partition coefficients). We propose performing sensitivity analyses to determine estimable parameters from the data. Depending on available tools, we can consider fitting the model to each animal separately then deriving representative parameters or pooling the animals. Alternatives like compartmental modelling to simplify the model structure or population approaches to simultaneously model several animals could also apply to PET data [11].

#### 2.1.2. Preclinical PK extrapolation from micro- to conventional study dose (Step B)

The second step bridges microdoses to therapeutic doses in the same preclinical species, addressing potential non-linearities from saturable ADME processes or target-mediated drug disposition [12]. Conventional PK data typically include sparse plasma and urine data, which can be repeatedly collected in larger species over a period of time. They may also include gallbladder bile and ex vivo tissue concentrations in studies involving animal sacrifice, providing longer observation windows to identify elimination routes and validate tissue distribution but losing the longitudinal aspect [12–14]. These findings can result from dose-escalation studies which provide PK information scaling with dose, and allow to determine toxic levels. NOAEL, representing the highest dose of a substance at which no significant toxicity is observed, is then commonly used to propose a safe dose range for first-in-human studies. Alternatively, target exposures can be derived from measuring and modelling response-related markers, including data from in vitro models [15], to determine the minimal anticipated biological effect level (MABEL) [16]. Major metabolites can be quantified in the samples using standard analytical techniques, providing information on metabolism rates. However, these data are only collected from a limited number of animals following ethical/practical constraints [17].

In this step, PK modelling (PBPK, pop-PK, etc.) comparing predictions from microdose PET studies to therapeutic exposures would address PK non-linearities, e.g, saturable CYP3A4/UGT metabolism, Pgp/BCRP transport, or target binding in TKIs [18]. Combining animal PET microdosing with sparse conventional PK data may inform PK modelling workflows to scale to first-in-human doses [15].

#### 2.1.3. Translation from preclinical to clinical PK using Human PET data (Step C)

Human PET microdosing offers valuable direct Phase 0 data in humans, mirroring preclinical PET to capture real-time tissue kinetics in the population of interest while benefiting from larger organ sizes for improved PET delineation and optimized sparse blood sampling for metabolites [15]. Despite short acquisition window, once human PET data are obtained, via dynamic imaging protocols tracking radiolabelled tracers in key organs (e.g., liver, kidney, brain) alongside venous blood draws, PK/PBPK models would enable targeted refinement. These models leverage built-in interspecies scaling (e.g., allometric exponents for body weight/flows, enzyme ontogeny adjustments) to bridge preclinical predictions to human data. The data can be analysed using the same modelling approaches as for preclinical data, benefitting from the same granularity as the animal PET data to explore species differences in organ distribution and fine-tune select parameters such as tissue partition coefficients, biliary/hepatic or transporter clearances [18]. They can also be combined via prior-informed uncertainty propagation for slower metabolism processes and metabolite distribution with preclinical data from the pharmacological doses, either by fixing specific parameters or by using Bayesian inference [19].

#### 2.1.4. Prediction of Human PK (Step D)

The models and optimised parameters from Steps A–C (microdose tissue distribution, non-linearity scaling, human data refinement) are then combined to predict human PK for the target exposure. Steps B (preclinical therapeutic scaling) and C (human microdose) are combined hierarchically: non-linearities identified in B (e.g., saturable clearances from sparse bile/plasma) inform priors for human PBPK fits in C, yielding a unified model for first-in-human dose prediction. Alternatively, predictions from partial paths (B or C) can be ensemble-averaged (e.g., model averaging weighted by uncertainty) to mitigate biases, such as higher animal–human extrapolation error from A+B alone or dose-scaling uncertainty from A+C (Phase 0 directly to Phase I) [20]. These steps build empirical priors for human dose-finding studies, e.g., preclinical predictions from previous frequentist or Bayesian analyses for virtual patient cohorts, enabling uncertainty propagation via MCMC sampling or sensitivity analysis on key parameters (partition coefficients, biliary clearance, etc.), thus supporting robust safety/efficacy margins (e.g., scaling from NOAEL or combining NOAEL and MABEL exposure targets). These simulations can be used to support dose selection for healthy volunteer dose-finding trial by evaluating simulated patient profiles across a range of candidate dose levels and applying constraints on exposure metrics such as C_max,_ C_trough,_ or AUC. This makes it possible to pre-identify doses that are likely to provide acceptable pharmacokinetic profiles in the target population while limiting the risk of excessive or insufficient exposure.

Note that this framework adapts to the available data: using only A+B would be associated with higher uncertainty on animal–human extrapolation, while using A+C (i.e., direct Phase 0 to Phase I transition) would incur higher uncertainty on dose scaling to therapeutic levels.

The full PBPK model could be too complex to integrate in a first-in-human study as is. As the PK study will rely on blood measurements, a further step could be to fit the simulated human profiles to a simplified compartmental model to obtain population parameter distributions as priors for PK dose-finding studies [21].

### 2.2. Dolutegravir case-study

We apply the framework to a case-study where we collected data for dolutegravir, an antiretroviral agent, from preclinical studies in the RHIVIERA project (https://rhiviera.com/project/imaging-drug-diffusion-and-viral-reservoirs-wp2/), mapping to Steps A and B from the framework with microdose PET and conventional-dose data respectively. In this example, dolutegravir is an approved drug with reported information about physico-chemical properties, high plasma protein binding, primary metabolism via UGT1A1 glucuronidation, and Pgp/BCRP-mediated transport pathways, as well as therapeutic PK exposure targets [22–24]. At this stage, these insights provide a strong basis for PBPK modelling and the framework evaluation, where the most important challenge was the unavailability of human PET data for step C, increasing uncertainty in NHP-to-human extrapolation.

The case-study data are summarised in detail in **Supplementary Information.** Briefly, PET microdosing data were collected in 4 NHPs receiving a single intravenous (IV) microdose of radiolabelled [^18^F]dolutegravir [25] alone or after an IV pre-injection of 2.5 mg/kg dolutegravir, (i.e., conventional dose) administered 1 hour earlier. n the analysis, we considered TACs obtained in selected organs as previously described [26]. PET data were converted to fraction of injected dose per organ volume (= *organ PET radioactivity(kBq/mL/) / injected radioactivity (kBq)* [27]. In arterial plasma samples, parent-specific data was quantified using radio-HPLC and converted to *fDTG*, the fraction of [^18^F]dolutegravir over total radiolabelled molecules (dolutegravir and principal metabolites M2/M6). Dolutegravir concentrations were quantified in fluids and tissues from NHPs administered a single conventional IV dose of dolutegravir alone or combined with tenofovir and emtricitabine, using LC-MS/MS as previously described [28,29].

#### 2.2.1. Framework application to dolutegravir

Framework application to dolutegravir is described in detail in **Supplementary Information.** Based on clinical relevance and drug distribution [22,30], data from blood (aorta on PET imaging or plasma concentrations), kidney, liver, urine (bladder), spleen, and plasma *fDTG* were used for PBPK model fitting across both microdosing and conventional-dose datasets. Measurements in other organs were used exclusively for model evaluation. A whole-body PBPK model was built using OSPSuite (PK-Sim/MoBi, version 12.1) [31]. All drug-related model parameters considered potentially estimable in this study are shown in **Supplementary Table S1**. Model parameters for estimation were selected from **Supplementary Table S1**, using one-at-a-time (OAT) sensitivity analyses. The model was fitted sequentially, first to PET microdosing data, then conventional-dose data. After each estimation, the model would be checked graphically if predictions aligned with the 2-fold deviation range. Finally, PK extrapolation from NHPs to humans was performed from the geometric mean parameters θ_C_ yielded from the final individual parameter estimates in NHPs after step B.

#### 2.2.2. Framework evaluation

To evaluate the ability of the framework to provide a reasonable description of human PK, we compared our PBPK model predictions with reported literature results. Concentration–time profiles were simulated in a population of 100 adults receiving the recommended dose of 50 mg oral dolutegravir qd for 5 days using θ_C_, mean parameters extrapolated from step B. We calculated the median and range of blood concentration–time profiles at steady-state, derived secondary PK parameters (e.g., AUC_0-24_, C_max_, C_trough_), For comparison we also computed the predictions obtained with the parameters extrapolated from model fitting to PET microdosing data (θ_P_) with our simulated concentration–time plots and secondary PK parameters (namely, AUC_0-24_, half-life, C_max_, T_max_, and C_trough_).

## 3. RESULTS

### 3.1. Dolutegravir PBPK modelling in NHPs using PET microdosing data

The full results for the sensitivity analysis to determine parameter identifiability are given in **Supplementary Figure S1**. Briefly, Vmax and Km were highly correlated for all enzymes in the model, and we chose to estimate only V_max_; parameters having no impact on PK parameters were set to default values (**Supplementary Table S1**). Finally, as some individuals did not have metabolite data, we set metabolite renal/biliary specific clearance (*CL_M_*), dolutegravir lipophilicity (*logP_DTG_*) and Vmax of transporters Pgp/BCRP to the same value in all subjects (*V_max, tr_*), and the same unbound fraction was set for dolutegravir and its metabolites (*fu*).

We then estimated the parameters simultaneously for all NHP PET data. **Table 1** shows the individual and geometric mean estimates for interstitial/plasma partition coefficients for kidney, liver, and spleen (*K_int:pl_*), *fu*, *CL_DTG_*, *CL_M_*, *V_max,tr_,* and *V_max,UGT_*. The geometric mean of *logP_DTG_*was 4.21, and the geometric mean *fu* of 0.89% was close to the initial value. Liver and kidney intracellular-to-plasma partition coefficients showed inter-individual variability, with *K_int:pl, Liv_* ranging from 10.78 to 15.60, and kidney *K_int:pl, Kid_* from 1.23 to 2.72.

**Table 1.** Model estimates using PET microdosing data.

| Parameter | P1 | P2 | P3 | P4 | Geometric mean |
| --- | --- | --- | --- | --- | --- |
| $\log P_{DTG}$ (Log Units) | 4.21 | | | | |
| $f_u$ (%) | 0.94 | 0.77 | 0.73 | 1.17 | 0.89 |
| $CL_{DTG}$ (1/min) | 2.78 | 2.63 | 1.10 | 3.97 | 2.38 |
| $CL_M$ (1/min) | 30.61 | | | | |
| $V_{max, tr}$ ($\mu\text{mol/l/min}$) | 176.17 | | | | |
| $V_{max, UGT}$ ($\mu\text{mol/l/min}$) | 540.74 | 409.00 | 1087.32 | 425.80 | 565.68 |
| $K_{int:pl, Kid}$ | 2.22 | 1.23 | 1.84 | 2.72 | 1.92 |
| $K_{int:pl, Liv}$ | 14.39 | 10.78 | 15.6 | 12.08 | 13.08 |
| $K_{int:pl, Spl}$ | 0.83 | 0.89 | 0.99 | 1.02 | 0.93 |

**Table 2.** Model estimates using preclinical concentration data. Starting values were set to estimates or geometric means of the corresponding individual parameters optimised with microdosing PET data. *, fixed to PET geometric means of metabolite specific clearance and interstitial/plasma partition coefficients for kidney, liver, and spleen (Kint:pl) after sensitivity analyses.

| Parameter | C1 | C2 | C3 | C4 | C5 | C6 | Geometric mean |
| --- | --- | --- | --- | --- | --- | --- | --- |
| $\log P_{DTG}$ (Log Units) | 3.90 | | | | | | |
| $f_u$ (%) | 0.35 | 0.61 | 0.65 | 0.66 | 0.65 | 0.71 | 0.59 |
| $CL_{DTG}$ (1/min) | 1.33 | | | | | | |
| $CL_M$ (1/min) | 30.61* | | | | | | |
| $V_{max, tr}$ ($\mu\text{mol/l/min}$) | 214.3 | | | | | | |
| $V_{max, UGT}$ ($\mu\text{mol/min/kg tissue}$) | 1669.8 | 988.3 | 406.1 | 495.1 | 544.1 | 503.3 | 670.5 |
| $K_{int:pl, Kid}$ | 1.92* | | | | | | |
| $K_{int:pl, Liv}$ | 13.08* | | | | | | |
| $K_{int:pl, Spl}$ | 0.93* | | | | | | |

The PBPK model adequately described radioactivity–time profiles following microdose administration of [^18^F]dolutegravir in all NHPs (**Figure 2**, **Supplementary Figure S2**). Individual predictions closely followed the observed data included in model fitting, namely *fDTG* in arterial plasma, and fraction of injected radioactivity in aorta, kidney, liver, urine (bladder), and spleen over 3 hours, reproducing closely both the early rise and decline phases, with most observations within 2-fold range (**Figure 2**). However, the model underpredicted liver C_max_ while overpredicting *fDTG* and late concentrations in kidney. In the organs not used for fitting, the model tended to overpredict brain concentrations, early bone uptake, late lung elimination, underpredicting peaks in heart, and poorly captured bile (gallbladder) profiles (**Figure 2**).

**Figure 2.**
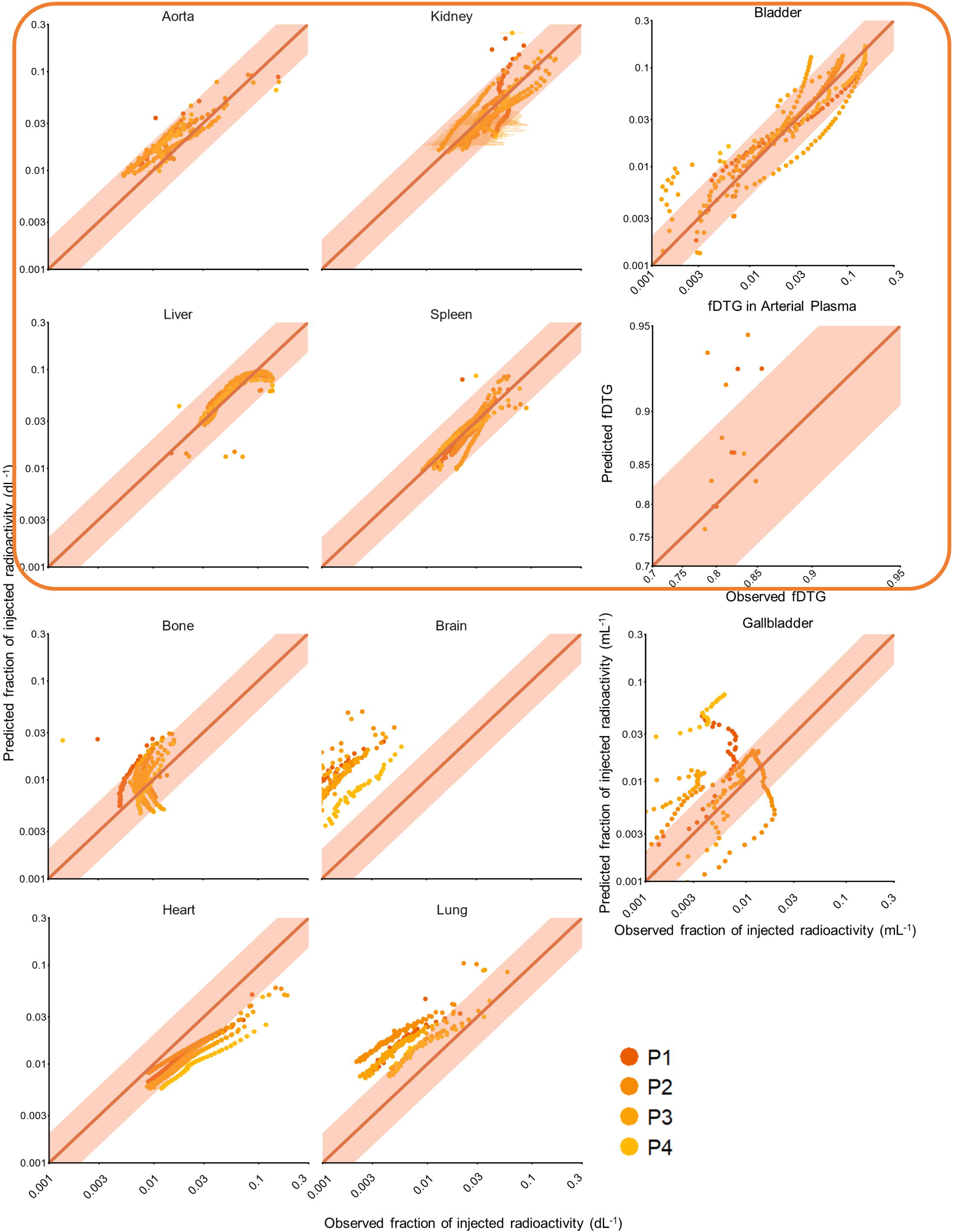
Predicted versus observed concentrations in NHPs receiving PET microdose of dolutegravir (P1-P4). Transparent ribbon: 2-fold deviation range; upper (framed): observed data included in model fitting; lower: observed data used only to compare with model prediction.

### 3.2. Modelling dolutegravir concentrations in NHPs receiving conventional dose

The sensitivity analysis (**Supplementary Figure S3**) for this stage suggested selecting *logP_DTG_*, *fu* of dolutegravir and metabolites, *CL_DTG_*, *V_max, tr_* and *V_max, UGT_* for re-estimation, with individual estimates for *fu* and *V_max, UGT_*, while CL_M_ and organ *K_int:pl_* were no longer considered estimable and fixed to PET estimates (**Table 1**). Comparing with parameter values from PET-derived data, conventional-dose estimates remained in the same order of magnitude as their PET-based priors. However, the therapeutic dose-based fit consistently yielded higher *CL_DTG_*and *V_max, UGT_*, with lower liver and kidney partition coefficients than the PET-derived set.

The model reproduced the overall magnitude of dolutegravir plasma and urine concentration–time profiles at 2.5 mg/kg IV (**Figure 3**). It also captured the kidney and spleen concentrations, with most measurements lying within the 2-fold deviation range, but overpredicted liver concentrations. Notably, in the organs not used for fitting, the model well captured heart concentrations. For bile, lung and brain, the model underestimated the observed values, with data points lying below the 2-fold deviation range (**Figure 3**).

**Figure 3.**
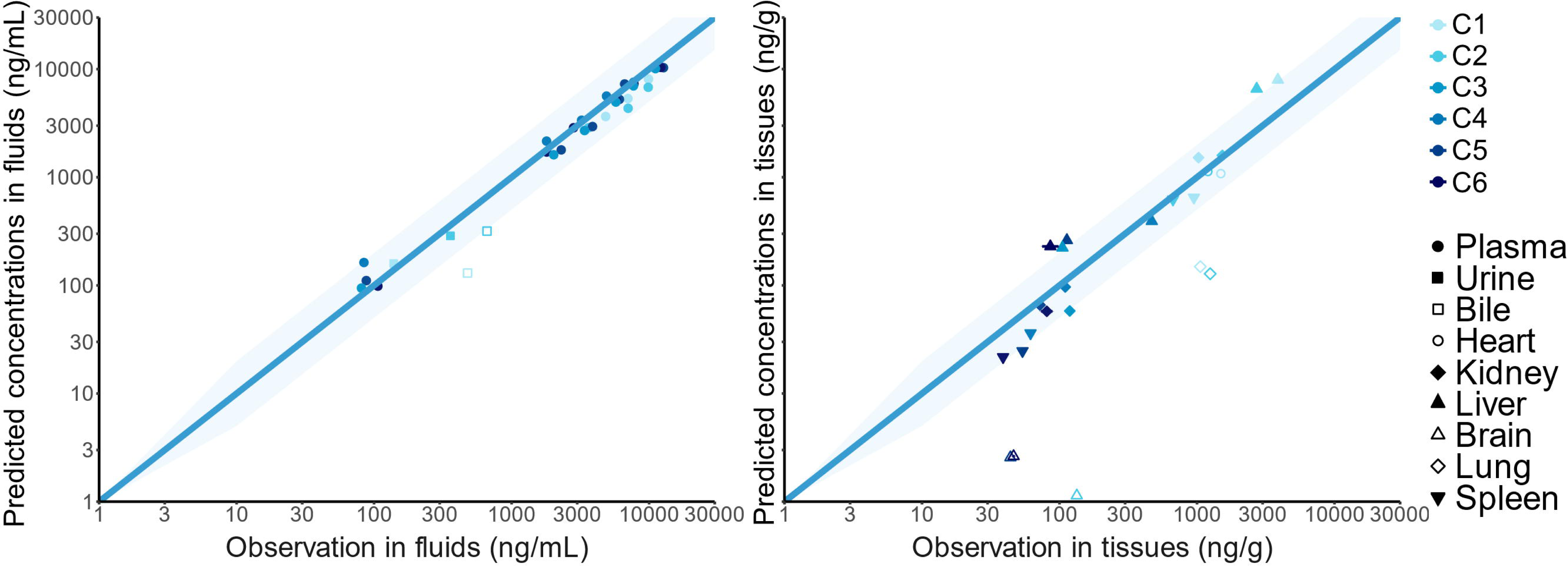
Predicted versus observed concentrations from fluids (left) and tissues (right) in NHPs receiving single injection of 2.5 mg/kg dolutegravir (C1-C6). Transparent ribbon: 2-fold deviation range; filled and empty symbols correspond to data included and not included in model fitting respectively.

### 3.3. Framework evaluation with healthy volunteers PK profiles

Five datasets from the dolutegravir arm of three phase I clinical studies meeting predefined criteria were extracted to evaluate model predictions of steady-state dolutegravir plasma pharmacokinetics in healthy volunteers. Study designs, dosing regimens, demographics, and sampling details for the included datasets are summarised in **Supplementary Table S2**. Datasets originated from Walimbwa et al. (2019 [33]; two cohorts of healthy volunteers), Johnson et al. (2014; two cohorts) [34], and Pene-Dumitrescu et al. (2021) [35].

Both NHP-derived parameter sets were applied to humans to simulate steady-state dolutegravir 50 mg once-daily dosing (**Figure 4**). Compared to literature reported data, the PET-only parameterisation (θ_P_) systematically underpredicted all metrics: AUC_0-24_ by 45%, C_max_ by 29%, C_trough_ by 82%, and t_1/2_ by 64% compared to dolutegravir reported reference values [22], thus under-predicting dolutegravir concentrations after peak. In contrast, the conventional-dose parameterisation (θ_C_) yielded predicted AUC_0-24_ and C_max_ aligned with the dolutegravir reported reference values. However, C_trough_ was 36% lower (but its prediction interval overlapped with 3 of the datasets) and t_1/2_ 45% shorter, leading to visible downward bias in the late portion of the curves (**Figure 4**, upper right).

**Figure 4.**
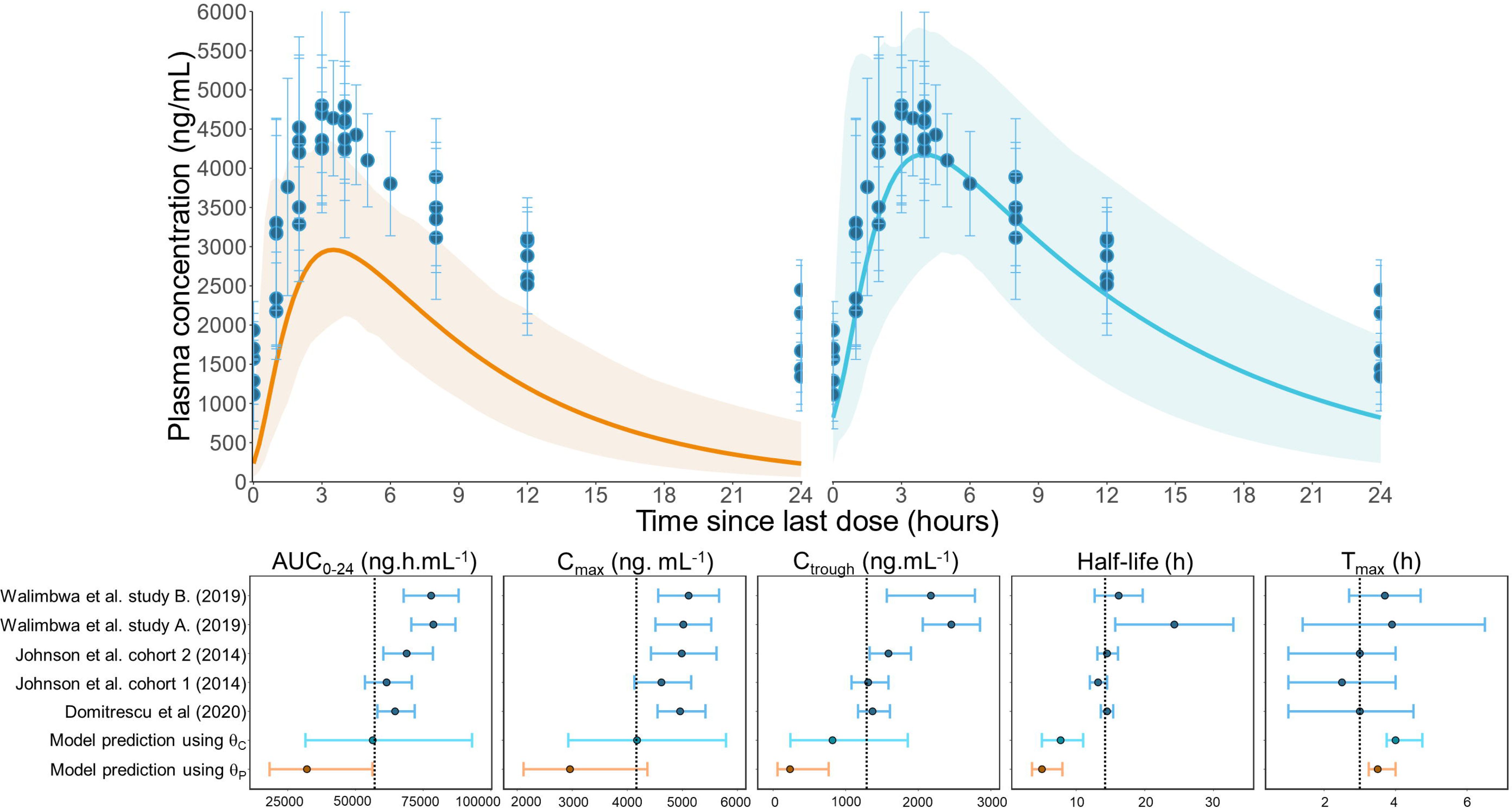
Dolutegravir predicted vs. reported pharmacokinetics at steady state in healthy volunteers using model estimates with preclinical PET microdosing and conventional PK data. Upper: Plasma predicted concentrations over time using θ_P_, parameter estimates from preclinical PET microdosing data, (left) and θ_C_, estimates refined sequentially from preclinical conventional PK data (right); lower: predicted secondary pharmacokinetic parameters; dotted vertical line: value reported by drug manufacturer. Results displayed as geometric means and 95% prediction intervals.

## 4. DISCUSSION

In this study, we designed a PET-informed PBPK framework that integrates microdose PET imaging data with therapeutic-dose concentrations to guide first-in-human dose selection. Applied to dolutegravir, a commonly prescribed antiretroviral agent for which we had PET and conventional-dose data in NHP, we showed that by leveraging microdosing and conventional-dose studies, the workflow could predict steady-state plasma concentrations in healthy volunteers within observed clinical range.

In the dolutegravir case study, extrapolation was achieved despite data limitations, including the short acquisition window due to radionuclide decay, a very small number of animals and sparse NHP sampling in the conventional-dose study. Compared to our full conceptual framework, we also lacked human PET microdosing data. Nonetheless, the PBPK model applied to PET data captured dolutegravir high plasma protein binding with *fu* estimated at 0.89%, consistent with known value between 0.5–2% [36–38]; *V_max, UGT_* of 565 µmol/min/kg was in the range expected for UGT1A1 substrates [24,39], while partition coefficients for liver, kidney and spleen (13.08, 1.92, and 0.93, respectively) aligned with reported ranges in the literature [40]. Model refinement from PET to therapeutic-dose parameters remained coherent, revealing dose-dependent nonlinearities. Sequential model fitting predicted higher *V_max,UGT_* with reduced *fu*, consistent with saturation emerging at 2.5 mg/kg IV. Most estimated parameters remained within the same order of magnitude. Inter-individual variability was high for coefficient partitions, which is plausible with short PET acquisition window, with more stable fraction of unbound dolutegravir across individuals related to drug elimination, which is better informed by longer conventional PK data.

PBPK modelling leverages physiological concepts enabling extrapolation across species, body size and administration regimens, making it particularly appealing in translational approaches. However, PBPK-based modelling generally relies only on plasma or local tissue PK and in vitro data [6,7,9,10,41–43]. Here, PET imaging provided high temporal resolution that enabled precise capture of initial distribution dynamics across organs and allowed to inform parameters of PBPK models that are usually collected from diverse in vivo and tissue models. Tissue-specific discrepancies emerged, with particularly poor capture of brain and bile profiles. This may likely result from multiple factors, including temporal mismatches between dynamic PET time-activity curves and static ex vivo measurements, pooled values used to develop the PBPK models in the modelling platforms such as PK-Sim and SimCYP, which consequently do not reflect biological variability, as well as dose-dependent saturation effects not apparent at microdose scale.

PET data limitations, besides the challenges mentioned in methods, include the inability to distinguish between parent drug and metabolites. Metabolite data, derived from the fraction of [^18^F]dolutegravir relative to total radiolabelled molecules in arterial plasma following PET microdosing, enabled initial partitioning of clearance between parent drug and metabolites, but was less frequently measured, limited to plasma and not available in all animals, adding uncertainty about absolute metabolite concentrations and tissue-specific distribution for parameter estimation. In the case of dolutegravir, most of the drug remained as parent drug over the early 3-hour acquisition window, but this could be an issue for rapidly metabolised drugs. The second step of the workflow was crucial in improving the estimates of terminal clearances, as seen when comparing the predictions of human PK profiles. In this study however, we had access to only 6 animals with sparse plasma concentrations and limited tissue samples. Re-estimating the parameters provided adequate description of plasma profiles, but some tissues were not well fitted, partly attributable to high inter-individual variability in tissue distribution evident from the dispersion in observed data, and methodological differences between PET fraction-of-dose metrics and LC-MS tissue concentrations.

In our dolutegravir case-study, human PET data was unavailable. In the framework, this step would be used to tailor the interspecies extrapolation to the target human population using PET dynamic data [2,44]. The framework could also accommodate several species to better identify differences in saturation across species or doses, especially since we couldn’t estimate both Km and Vmax in the associated metabolism processes [45,46]. Human PET would also provide prediction of tissue exposure, relevant to set safe starting doses before proceeding to first-in-human therapeutic dosing and vulnerable population assessment. However, these studies could be limited by ethical or practical constraints, e.g., the number of inclusion or duration of the experiment, as well as interspecies uncertainties involving protein abundance or activity.

In our application, we were limited by the methodological approaches available in the current implementations of PBPK platforms. Local sensitivity analysis, as implemented in PK-Sim/MoBi [32,47], was employed for computational efficiency, but examines parameters individually, potentially missing correlated interactions (e.g., between *V_max,UGT_* and *K_int:pl, Liv_*). Global methods [32,40,47] would quantify interactions but require substantial computational resources. Parameter identifiability remains challenging, concerning either model structure or data-limited issues [48], which is the case here where sparse multi-organ data likely confound parameter trade-offs despite OAT sensitivity screening. Moreover, our estimation assumed uniform residual error across tissues, which is problematic given analytical method and tissue discrepancies [49]. Our study yielded only individual fits, which lacks interindividual structure, which should be subsequently accommodated by Bayesian MCMC [6,7] or SAEM [49]. Moreover, parameter uncertainty was unavailable and therefore not incorporated into the simulation. If parameter uncertainty becomes available, a probabilistic sensitivity analysis could be incorporated within this framework to inform Bayesian prior distributions for prospective PK analyses, not only for the mean but also for the variance, as an indicator for the information amount.

Whole-body PBPK modelling was essential for this analysis, as plasma-only population PK approaches would likely be inadequate, requiring a substantially larger sample size and rich sampling to ensure stable parameter estimation [11], given the challenge of bridging microdose to therapeutic exposures. The PBPK framework, by incorporating physiological determinants, such as organ blood flows and PET-derived tissue–plasma partition coefficients, enabled mechanistic scaling across dose levels. PET data offers unique value for PBPK beyond in vitro or plasma data only, i.e., whole-body, dynamic data across various organs in NHPs in vivo, with high temporal resolution capturing initial distribution unavailable from sparse LC-MS. However, this mechanistic detail introduced additional complexity, involving more than 50 parameters, many fixed from literature or auxiliary data [48,49], and tissue concentration prediction quality remained variable, suggesting that the perfusion-limited assumption or unmodeled binding saturation may have constrained the model predictive accuracy.

In conclusion, this proof-of-concept study demonstrates the feasibility of a sequential PET microdosing-to-therapeutic dose PBPK workflow for NHP-to-human pharmacokinetic extrapolation, leveraging dynamic whole-body imaging data with complementary tissue sampling. By exploiting imaging-derived organ time–activity curves and classic concentration data in NHPs, the model achieved acceptable prediction of human steady-state exposure at a therapeutic dose, thereby supporting the initial objective of assessing phase 0-PBPK strategies for cross-species extrapolation. In a prospective setting, this framework combines PET microdosing in animals to quantitatively inform initial PBPK model construction, therapeutic-dose extrapolation to explore non-linearities, and human microdosing PET studies, which are explicitly supported by FDA/EMA phase 0 guidance and ICH M3(R2) thanks to their substantial safety margins [2]. Target-specific PET imaging (e.g., lymph nodes, viral sanctuaries) [25] could also help to determine the target exposure. Further methodological refinement, particularly with respect to tissue processes and estimation methods, should be explored before such PET-informed PBPK workflows can be routinely deployed for first-in-human dose selection and broader translational decision-making.

## Declarations

### Data Availability

The datasets generated during and/or analysed during the current study are available from the corresponding author on reasonable request.

### Ethics Approval

The study protocols were approved by the French regulatory authority (APAFIS_2453-2015102713323361 for the THE907 study and APAFIS#28763-2020122115083093 for the THE2001 study). The preclinical PET study was approved by the local ethical committee (CETEA n°44) and the French Ministry of Education and Research (APAFIS #40586-2023013115557146 v1).

### Consent to Participate

Not applicable.

### Consent for Publication

Not applicable.

### Code Availability

Not applicable.

### Author Contributions

B.T.N., E.C., and M.U. wrote the manuscript; N.T. and E.C. designed the research; B.T.N, A.B.-T., N.T., S.G., and F.C. performed the research; B.T.N, E.C, and M.U analysed the data; A.B.-T., S.G., F.C., T.N., R.L.G., O.L., and N.T. contributed new reagents/analytical tools.

## Acknowledgements

Perplexity was used for language editing to improve readability and coding trackability for figure generation; authors reviewed and take full responsibility for the final content.

---

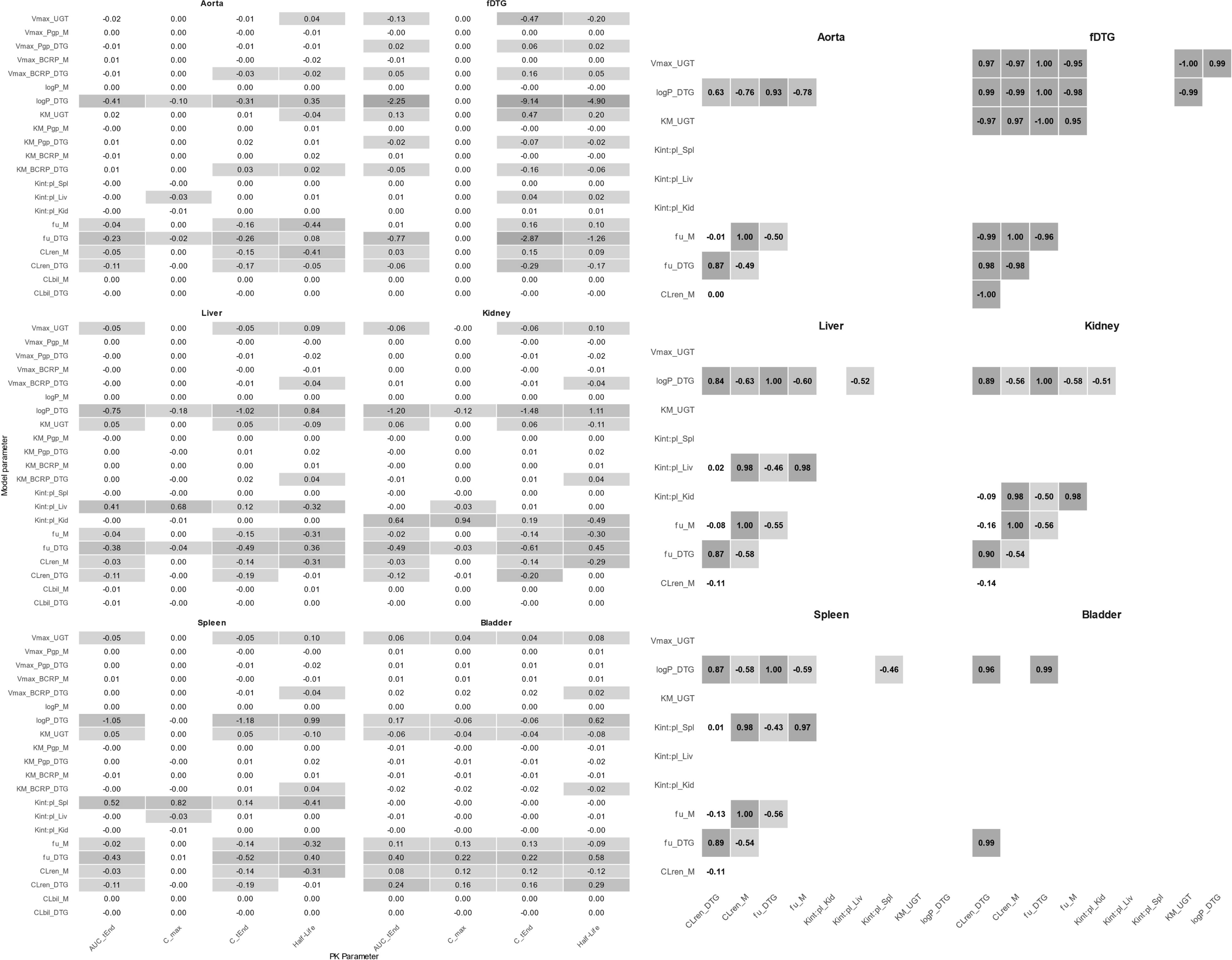

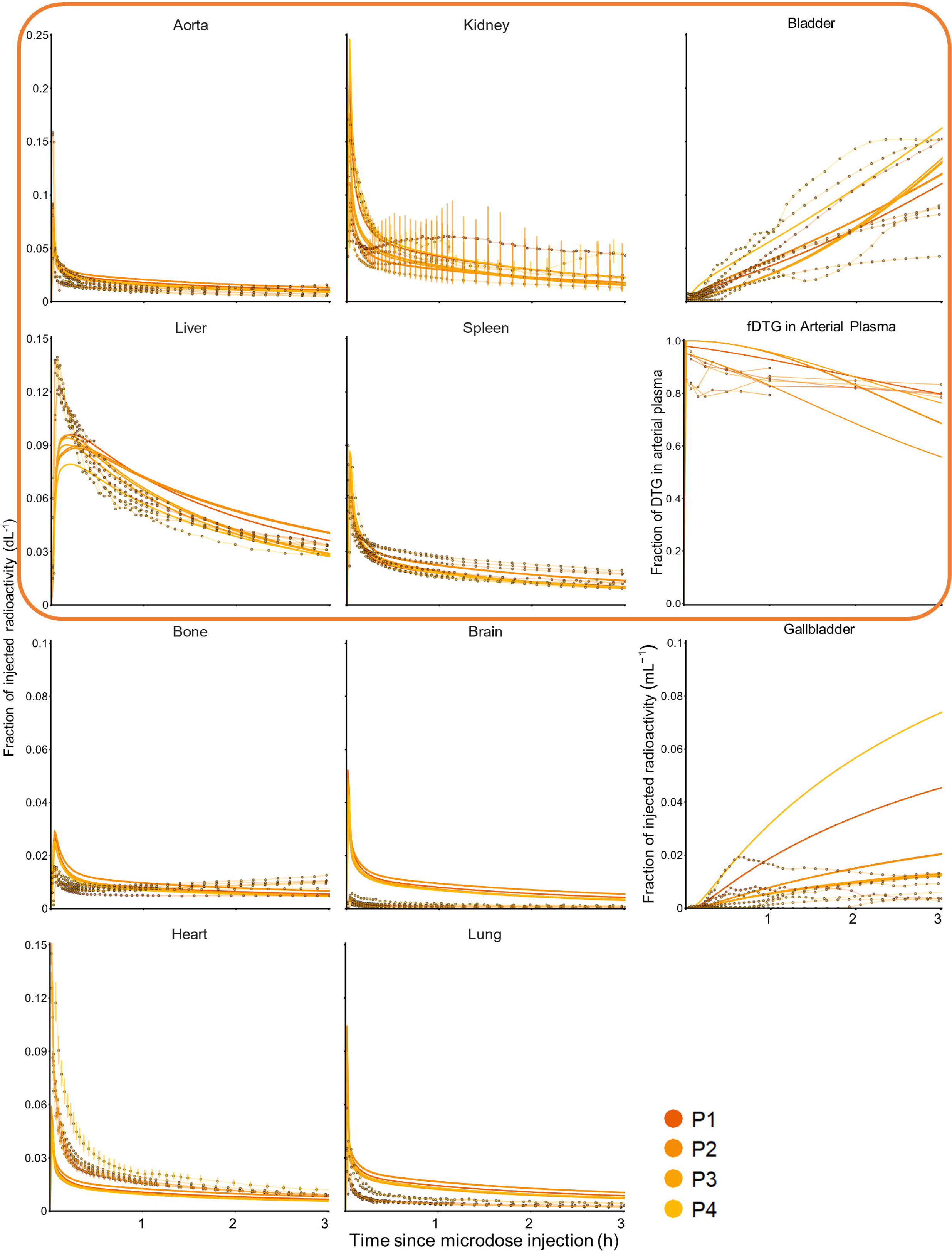

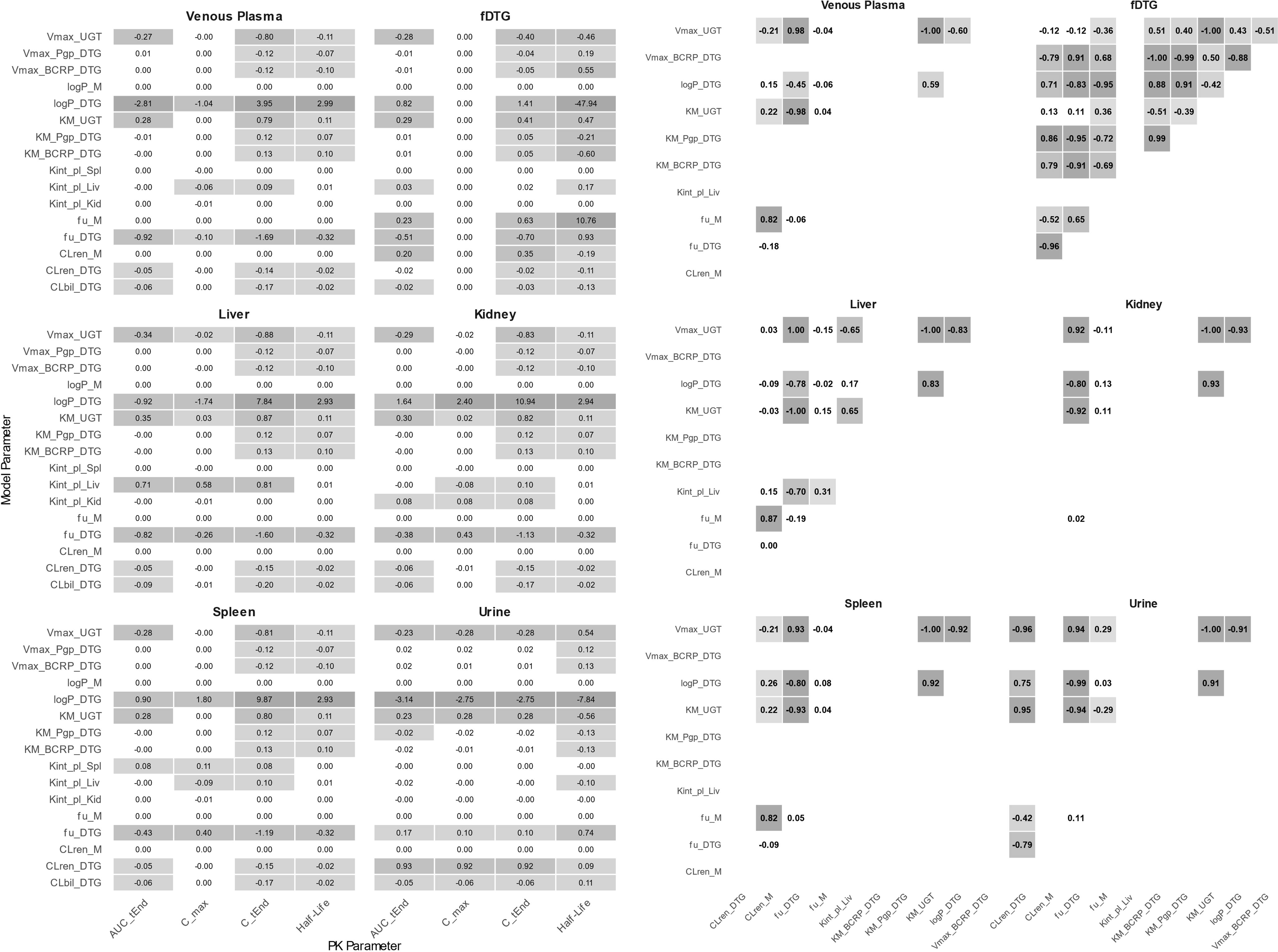

