## Supplementary Information for "A novel workflow integrating whole-body PET microdosing data and therapeutic-dose pharmacokinetics across species to inform first-in-human dose selection"

B.T.Nguyen, A.Barrail-Tran, M.Ursino, S.Goutal, F.Caillé, T.Naninck, R.Le Grand, O.Lambotte, N.Tournier, E.Comets

A - Dolutegravir case-study

A.1 Data

#### PET imaging data in NHPs receiving dolutegravir single microdose

PET microdosing data were collected in 4 male cynomolgus macaques (8.8 ± 0.8 kg, 7.5 ± 2.5 years. Two animals received a single intravenous (IV) microdose (0.02-1 µg/kg) of radiolabelled dolutegravir, 1 animal received a microdose preceded by an IV injection of 2.5 mg/kg dolutegravir, (i.e., conventional dose) administered 1 hour earlier, with the experiment repeated on 2 occasions. The last animal received a microdose alone on 2 occasions and after pre‑injection on 1 occasion.

Radiolabelled [^18^F]dolutegravir was produced onsite using the TRACERlab® FX N Pro module (GE Healthcare, Buc, France) as previously described (Tisseraud et al., 2022). Whole-body distribution of [^18^F]dolutegravir was assessed using a Siemens Biograph PET-CT scanner (Siemens Healthcare, TN, USA) under standard anaesthesia and monitoring procedures (Auvity et al., 2021). PET acquisition started with the IV bolus injection of [^18^F]dolutegravir (194.5 ± 36.0 MBq/kg, 0.40 ± 0.23 µg/kg). Dynamic acquisitions consisted of repeated whole-body PET images (3-bed step) during 180 min on a Siemens Biograph PET-CT scanner (Siemens Healthcare, USA). Dynamic PET images were reconstructed using standard PSF-OSEM-3D algorithms, with corrections for radioactive decay, scatter, attenuation, and detector inhomogeneity. PET image analysis was performed using PMOD software version 4.3 (PMOD Technologies Ltd., Zurich, Switzerland). TACs were obtained in selected organs including aorta, kidneys, liver, gallbladder, bladder, brain, bone, heart, lung, and spleen. For each organ and time, PET radioactivity was converted to fraction of injected dose per organ volume (

$$= organ PET radioactivity (kBq/mL) / injected radioactivity (kBq)$$

) (Schluep et al., 2009).

In arterial plasma samples, parent-specific data was quantified at 0, 5, 15, 30, 60, 120, and 180 minutes post-microdose using radio-HPLC and converted to *fDTG*, the fraction of parent dolutegravir over total radiolabelled molecules (dolutegravir and principal metabolites M2/M6).

#### Dolutegravir concentrations in NHPs administered a conventional dose

Preclinical studies were performed in the RHIVIERA project to characterise whole-body distribution of dolutegravir in NHPs. Two female NHPs from THE2001 received a single IV injection of 2.5 mg/kg dolutegravir alone, in which dolutegravir and metabolites (M2/M6) concentrations were measured in venous plasma at 1h, 2h, 3h post-injection, as well as in urine, bile, and tissues including liver, kidney, heart, lung, and spleen, at euthanasia 3 hours post-injection. In a second study (THE1907) (Gelé et al., 2024), 4 male NHPs receiving a single injection of 2.5 mg/kg dolutegravir coadministered with tenofovir and emtricitabine, in which dolutegravir concentrations were measured in venous plasma, urine, and bile at 1h, 2h, 4h, 6h, 24h post-injection, and in tissues (namely, kidney, liver, brain, heart, lung, and spleen) at euthanasia at 24h post-injection. Dolutegravir quantification in fluids and tissues was previously described using LC-MS/MS (Gelé et al., 2024; Gouget et al., 2020).

**A.2 Methods**

#### PBPK model

Based on clinical relevance and drug distribution (Bevers et al., 2023; Seaton, 2013), data from blood (aorta on PET imaging or plasma concentrations), kidney, liver, urine (bladder), spleen, and plasma fraction of dolutegravir were used for PBPK model fitting across both microdosing and therapeutic-dose datasets. Brain, heart, lung, bone, and gallbladder measurements were used exclusively for model evaluation.

A whole‑body PBPK model of dolutegravir was built using OSPSuite (PK-Sim/MoBi, version 12.1) (Willmann et al., 2003). The structural model included standard PK‑Sim organs (arterial and venous blood, lung, liver, kidney, spleen, heart, brain, adipose tissue, muscle, gut, etc.), each divided into vascular, interstitial, and intracellular spaces according to the PK‑Sim small‑molecule template with perfusion‑limited distribution. Physicochemical properties of dolutegravir (molecular weight, lipophilicity, pKa, plasma protein binding, and the blood‑to‑plasma concentration ratio) were used to determine tissue distribution and permeability. The model included known PK properties of dolutegravir, including the strong binding of dolutegravir to plasma albumin, and hepatic metabolism through glucuronidation by UGT1A1 to form metabolites M2 and M6. We added active efflux processes mediated by P‑glycoprotein (Pgp) and breast cancer resistance protein (BCRP) at the relevant biological membranes, and accounted for both biliary and renal clearance of unchanged dolutegravir and its glucuronide metabolites.

Metabolites were assumed to share the same distribution model as dolutegravir, with distinct renal and biliary clearance terms, implemented as first‑order clearances from liver and kidney tissue. Physiological parameters (organ volumes, blood flows, glomerular filtration rate) for NHPs and humans were taken from the PK‑Sim species libraries and scaled to individual body weight and demographic characteristics.

All enzyme‑ and transporter‑mediated processes were assumed to follow Michaelis‑Menten kinetics:

$$v=V_{max}\times[S]/(K_{M}+[S])$$

, where *v* is the velocity of the enzymatic reaction or transport, and *V*max (μmol/min/μmol protein) is the maximum rate of the enzymatic reaction or transport, [*S*] is the substrate concentration (μM), and *K_M_* is the Michaelis‑Menten constant (μM). All drug-related model parameters considered potentially estimable in this study are shown in **Supplementary Table S1**, while other physiology-specific parameters were fixed to PK-Sim/MoBi default values. Partition coefficients and cellular permeabilities would then be calculated accordingly from these estimates with the PK-Sim Standard built-in method (Willmann et al., 2003).

#### Parameter Selection for Estimation

To select from **Supplementary Table S1** which model parameters to estimate, based on the available data included for model estimation, we performed PK-Sim/MoBi built-in one-at-a-time (OAT) sensitivity analyses for each individual organ data selected for the model fitting, by evaluating how variation in a parameter influenced PK parameters following drug administration, while keeping all the other parameters unchanged (Najjar et al., 2024). A model parameter was selected for estimation, if varying its value by 100% changed at least one PK parameter involving the organs included in model fitting by at least 20% (Najjar et al., 2024). When two such parameters corresponded to two PK parameter changes that were highly correlated (ρ>0.95), we fixed one of them to the reported value in the literature (e.g., fixing Km while optimising V_max_), or set it to the value of another similar model parameter (e.g., metabolite parameters set to dolutegravir value). If a parameter was considered estimable only in some individuals (e.g., metabolite parameters), we estimated a single shared parameter value applied to all individuals.

#### Sequential Parameter Estimation

A sequential process was performed to estimate individual parameters, where estimates obtained from the previous steps were used as initial value for the next optimisation. First, we fitted the model to PET data from all NHPs receiving an IV microdose of dolutegravir, summarising individual estimates as geometric means when appropriate. Next, we fitted the model to concentration data from NHPs following a single dose of 2.5 mg/kg, with initial values set to previous estimates. Of note, with little metabolite-related information particularly in preclinical concentration data due to sparse sampling in very few individuals, model parameters concerning metabolites, namely renal/biliary specific clearances, were fixed to estimates obtained with PET data. Also, for each optimisation step of the sequential estimation, the sensitivity analysis described above was performed again to determine which parameters could be estimated at each stage.

Parameters were estimated by minimizing the additive log-scaling residual sum of squares between observed data and model predictions. Data used for estimation were weighted proportionally to the number of available observations per individual and per data type (e.g., PET time-activity curves, plasma/tissue concentrations), with all individuals pooled at each estimation stage.

#### Model assessment

After each estimation, the model would be assessed graphically by checking if predictions aligned with the 2-fold deviation range, i.e., 50%-200% compared to observed data (on logit scale for *fDTG* data and linear scale for others).

*Model prediction and extrapolation to human PK*

In the final step, the PBPK model was extrapolated from NHP to humans using PK-Sim's built-in scaling relations, including allometric scaling for physiological parameters (e.g., organ volumes, blood flows), species-specific enzyme and transporter abundances, and activity distributions for UGT1A1-mediated metabolism and Pgp/BCRP efflux. We extrapolated the human values from the geometric mean of the final individual parameter estimates in NHPs fitted to PET microdosing and conventional-dose data after step A (θ_P_) and B (θ_C_) respectively, to simulate concentration versus time profiles in humans. Inter-individual variability was introduced via biological covariates, namely, body weight, height, age, and sex in the target population, which were simulated using the default PK-Sim distribution for healthy Caucasian adults. To predict the PK after oral doses, which is the formulation used in humans, we added the default PK-Sim build-in model of oral absorption for dissolved formulation, including fixed intestinal paracellular permeability

A.3 Results

The main results are described in the manuscript, and we detail here some additional results about sensitivity analyses to select model parameters for each estimation stage. The full results for the sensitivity analysis to determine parameter identifiability are given in **Supplementary Figure S1** and **Supplementary Figure S3,** respectively for estimation with PET microdosing and conventional-dose data. We tested by organ the impact of varying model parameters on pharmacokinetic (PK) parameters, as shown on the left panel. Only model parameters associated with at least 2 impacts beyond 0.2 across individuals and PK parameters would be tested for correlations between impacts of model parameters across individuals and PK parameters, as shown on the right panel**.** Empty cases imply that at least one model parameter of the pair unsatisfied the condition for correlation test.

B - Framework evaluation

B.1 Methods

The current recommended dosage for dolutegravir is 50 mg once daily (qd), and published PK data could be found for this regimen. We therefore systematically searched PubMed and Google Scholar for phase I studies with the following characteristics: (i) administration of dolutegravir 50 mg qd as monotherapy (most frequently investigated dosing regimens in phase I trials), (ii) rich PK sampling over at least 24 h at steady-state (after ≥5 days of dosing), and (iii) availability of extractable concentration-time profiles. Search terms included "dolutegravir steady-state pharmacokinetics healthy volunteers 50 mg" and "dolutegravir phase I PK monotherapy", published until January 2026. Plasma profiles displayed as population geometric means (± standard deviation) were digitised using WebPlotDigitizer (<https://automeris.io/WebPlotDigitizer>) and expressed in ng/mL.

We simulated a virtual population of 100 adults receiving the recommended dosage regimen of 50 mg qd oral dolutegravir for 5 days after a fat meal of 600 kcal reflecting the conditions of the published studies in healthy volunteers, using the physiological variability distributions built-in PK-Sim. The simulated population consisted in white American healthy volunteers, 50% male, aged 20 to 80 years, weighting 40 to 100 kg, with heights from 150 to 190 cm. We used PK-Sim built-in formulas to extrapolate to humans and adjust the model parameters to individual covariates. We predicted the median and range of blood concentration profiles at steady-state and derived secondary PK parameters (e.g., AUC_0-24_, C_max_, C_trough_). To assess the different components of the framework, we compared the predictions with those obtained with θ_P_ and θ_C_ by superimposing the prediction intervals with the confidence intervals from the observed data as reported in the published studies.

B.2 Results

**Supplementary Table S2** shows the studies extracted through the literature search, providing PK data and summary statistics for AUC and C_max_ of dolutegravir at steady state in patients receiving 50mg daily. The table briefly describes their characteristics, population and sampling schedules.

List of Supplementary Tables

Supplementary Table S1. Parameters used in the PBPK model for dolutegravir.

| **Parameter** | **Abbreviation** | **Unit** | **Start Value** | **Source** |
| --- | --- | --- | --- | --- |
| Lipophilicity of dolutegravir | *logP_DTG_* | Log Units | 2.20 | PubChem (PubChem, n.d.) |
| Lipophilicity of metabolites | *logP_M_* | Log Units | -0.88 | PubChem (PubChem, n.d.) |
| Fraction unbound of dolutegravir and metabolites | *fu* | % | 1.10 | PubChem (PubChem, n.d.) |
| Dolutegravir renal/biliary specific clearance | *CL_DTG_* | 1/min | 1.00 | PK-Sim Default |
| Metabolite renal/biliary specific clearance | *CL_M_* | 1/min | 1.00 | PK-Sim Default |
| K_M_ of Pgp and BCRP | *K_M, tr_* | µmol/l | 673.0* | Korzekwa and Nagar (2014) (Korzekwa & Nagar, 2014) |
| V_max_ of Pgp and BCRP | *V_max, tr_* | µmol/l/min | 1000.0* | Korzekwa and Nagar (2014) (Korzekwa & Nagar, 2014) |
| K_M_ of UGT1A1 | *K_M, UGT_* | µmol/l | 149.0 | Reese et al. (2013) (Reese et al., 2013) |
| V_max_ of UGT1A1 | *V_max, UGT_* | µmol/min/kg tissue | 409.0 | Reese et al. (2013) (Reese et al., 2013) |
| Kidney partition coefficient (interstitial/plasma) | *K_int:pl, Kid_* |  | 1.00 | PK-Sim Default |
| Liver partition coefficient (interstitial/plasma) | *K_int:pl, Liv_* |  | 1.00 | PK-Sim Default |
| Spleen partition coefficient (interstitial/plasma) | *K_int:pl, Spl_* |  | 1.00 | PK-Sim Default |
| Intestinal paracellular permeability (cm/s) | *P_intestinal_* | cm/s | 2.37×10^4^ | Reddy et al. (2021) (Reddy et al., 2021) |

Initial value for KM and Vmax of Pgp and BCRP, denoted with an *, were set to geometric mean of reported values in the source reference.

Supplementary Table S2. Clinical dolutegravir data used for PBPK model assessment.

| **Study** | **Design & Study Arm** | **Population (n)** | **Dolutegravir Dosing** | **Food Condition** | **PK Sampling Schedule** |
| --- | --- | --- | --- | --- | --- |
| **Walimbwa et al. (2019), Study A** | 2-way crossover study of drug-drug interaction with artemether-lumefantrine | 14 male healthy volunteers; median (IQR) age 29 years (21‑32), weight 55.5 kg (54‑64), BMI 21 kg/m² (17‑23) | 50 mg once daily for 6 days | Moderate-fat meal | On Day 6: 0 h (predose), 1, 2, 3, 4, 8, 12, 24 h postdose |
| **Walimbwa et al. (2019), Study B** | Drug-drug interaction study with artesunate-amodiaquine; parallel design | 13 male healthy volunteers; median age 30.5 years (23.5‑34), weight 60 kg (58‑68), BMI 20.5 kg/m² (19‑24.5) | 50 mg once daily for 7 days | Moderate-fat meal | On Day 7: 0 h (predose), 1, 2, 3, 4, 8, 12, 24 h postdose |
| **Johnson et al. (2014), Cohort 1** | Drug-drug interaction study with boceprevir; one-way study; dolutegravir monotherapy (Period 1) | 15 healthy volunteers; mean ± SD age 42.5 ± 17 years, weight 74 ± 13 kg, BMI 26 ± 3 kg/m²; 9 male, 11 Caucasian | 50 mg once daily for 5 days (Period 1) | Moderate-fat meal (30 g fat, 600 kcal) | 0 h (predose), 1, 2, 3, 4, 8, 12, 24 h postdose (Day 5) |
| **Johnson et al. (2014), Cohort 2** | Drug-drug interaction study with telaprevir; one-way study; dolutegravir monotherapy (Period 1) | 16 healthy volunteers; mean ± SD age 40 ± 15 years, weight 74 ± 14 kg, BMI 26 ± 3.5 kg/m²; 9 male, 11 Caucasians | 50 mg once daily for 5 days (Period 1) | Moderate-fat meal (30 g fat, 600 kcal) | 0 h (predose), 1, 2, 3, 4, 8, 12, 24 h postdose (Day 5) |
| **Pene-Dumitrescu et al. (2021)** | Drug-drug interaction study with GSK3640254; fixed-sequence, 3-period; dolutegravir monotherapy (Period 1) | 16 healthy adults (30 screened, 16 enrolled and completed); mean ± SD age 37 ± 11 years, weight 78 ± 10 kg, height 170 ± 6 cm, BMI 27 ± 2.8 kg/m²; 15 males; 11 White, 3 Black/African American, 2 of Asian heritage | 50 mg once daily for 5 days (Period 1) | Moderate-fat meal (30 min before dosing; 30 g fat) | 0 h (predose) Days 2‑5; postdose: 1, 1.5, 2, 3, 3.5, 4, 4.5, 5, 6, 8, 12, 24, 48, 72 h after Day 5 dose |

List of Supplementary Figures

Supplementary Figure S1. Sensitivity analyses to select parameter for model fitting to PET data.

Left: Impact of varying model parameters on pharmacokinetic (PK) parameters tested by organ in individual P3; right: correlations between those impacts of model parameters across individuals and PK parameters; only model parameters associated with at least 2 impacts beyond 0.2 across individuals and PK parameters would be tested for correlations; empty cases (right): at least one model parameter of the pair unsatisfied the condition for correlation test.

Supplementary Figure S2. Individual prediction for PET microdosing data.

Points: observations; vertical line: standard deviation (kidney and heart data only); bold: predictions; upper (framed): observed data included in model fitting; lower: observed data used only to compare with model prediction.

Supplementary Figure S3. Sensitivity analyses to select parameters for model fitting to conventional-dose data.

Left: Impact of varying model parameters on pharmacokinetic (PK) parameters tested by organ in individual C1; right: correlations between those impacts of model parameters across individuals and PK parameters; only model parameters associated with at least 2 impacts beyond 0.2 across individuals and PK parameters would be tested for correlations; empty cases (right): at least one model parameter of the pair unsatisfied the condition for correlation test.
